# Cardiovascular Hemodynamics in Mice with Tumor Necrosis Factor Receptor - Associated Factor 2 Mediated Cytoprotection in the Heart

**DOI:** 10.1101/2022.09.03.506447

**Authors:** Andrea G. Marshall, Kit Neikirk, Zer Vue, Heather K. Beasley, Edgar Garza-Lopez, Larry Vang, Taylor Barongan, Zoe Evans, Amber Crabtree, Elsie Spencer, Josephs Anudokem, Remy Parker, Jamaine Davis, Dominique Stephens, Steven Damo, Thuy T. Pham, Jose A. Gomez, Vernat Exil, Dao-fu Dai, Sandra Murray, Mark L. Entman, George E Taffet, Antentor O. Hinton, Anilkumar K. Reddy

**Author notes:** **Corresponding Authors Anilkumar K. Reddy, Ph.D.**, Department of Medicine, Baylor College of Medicine, One Baylor Plaza, MS: BCM285, Houston, TX 77030, USA., **Antentor O. Hinton, Jr, Ph.D.**, Department of Molecular Physiology and Biophysics, Vanderbilt School of Medicine Basic Sciences, The Vanderbilt Diabetes Research and Training Center, 750 Robinson Research Building, 2200 Pierce Ave, Nashville, TN 37232-0615.

## Abstract

Many studies in mice have demonstrated that cardiac-specific innate immune signaling pathways can be reprogrammed to modulate inflammation in response to myocardial injury and improve outcomes. While the echocardiography standard parameters of left ventricular (LV) ejection fraction, fractional shortening, and end-diastolic diameter, and others, are used to assess cardiac function, their dependency on loading conditions somewhat limit their utility in completely reflecting the contractile function and global cardiovascular efficiency of the heart. A true measure of global cardiovascular efficiency should include of the interaction between the ventricle and the aorta (ventriculo-vascular coupling, VVC) as well as measures of aortic impedance and pulse wave velocity. We measured cardiac Doppler velocities, blood pressures, along with VVC, aortic impedance, and pulse wave velocity to evaluate global cardiac function in mouse model of cardiac-restricted low levels TRAF2 overexpression that conferred cytoprotection in the heart. While previous studies reported that response to myocardial infraction and reperfusion was improved in the TRAF2 overexpressed mice, we found that TRAF2 mice had significantly lower cardiac systolic velocities and accelerations, diastolic atrial velocity, lower aortic pressures and rate-pressure product, lower LV contractility and relaxation, and lower stroke work when compared to littermate control mice. Also, we found significantly longer aortic ejection time, isovolumic contraction and relaxation times, and significantly higher mitral early/atrial ratio, myocardial performance index, and ventricular vascular coupling in the TRAF2 overexpression mice compared to their littermate controls. We found no significant differences in the aortic impedance and pulse wave velocity. While the reported tolerance to ischemic insults in TRAF2 overexpression mice may suggest enhanced cardiac reserve, our results indicate a diminished cardiac function in these mice.

## Introduction

Tumor Necrosis Factor (TNF) Receptor Associated Factor 2 (TRAF2), an E3 Ubiquitin-protein ligase, interacts with signaling molecules responsible for cell death, cellular stress response, activation of nuclear factor-*k*B and JNK (Bradley and Pober, 2001; Wajant and Scheurich, 2001) and critical pathways for cellular senescence and regeneration (Yarza et al., 2016). The TRAF2 pathway is activated by tumor necrosis factor and recruits for apoptosis inhibitor proteins, cIAP1/2 (Gough and Myles, 2020). Tumor necrosis factor can protect against ischemic-induced cardiomyocyte death (Miao et al., 2020). In contrast, its knockout is less cardioprotective (Benjafield et al., 2001; Higuchi et al., 2004). In cancer studies it was shown that TRAF2 can act as both a tumor promotor and suppressor (Siegmund et al., 2022). Given such juxtaposing roles, it is not surprising that mixed outcomes have been reported on the role of TRAF2 as a potential downstream mediator of cardiac protection in heart failure. TNF is cytoprotective in the heart via TRAF2 mediated activation of NF-kB (Burchfield et al., 2010), TRAF2 mitochondrial localization is cytoprotective, and may also contribute to TRAF2-mediated mitophagy (Yang et al., 2015), and that TRAF2 mediates myocardial survival by suppressing apoptosis and necroptosis (Guo et al., 2017). These roles make TRAFs important targets in many inflammatory and autoimmune diseases (So, 2022). However, other studies have suggested that cardiac function was improved after myocardial infarction in absence of NF-kB p50 subunit (Frantz et al., 2006), that suppressing NF-kB with oligonucleotide decoys reduces ischemia related myocardial injury (Tranter et al., 2012), and that TRAF2 mediates enhanced cardiac hypertrophy via activation of AKT/GSK3β signaling (Huang et al., 2014). Regardless, overexpression of TRAF2 was found to have cardioprotective role in myocardial injury and perhaps can be novel therapeutic strategy in cardiomyopathy (Ma et al., 2022).

Most studies in mice use echocardiography measurements of LV ejection fraction (EF), fractional shortening (FS), and end-diastolic diameter (EDD), among others, to assess cardiac function much akin to those used clinical studies (Tona et al., 2021). But EF, FS, or EDD are dependent on loading conditions and not completely reflective of the contractile function of the heart (Park et al., 2018). Therefore, it is important include the interaction between ventricle and the aorta (also known as, ventricular-vascular coupling, VVC) to evaluate the performance of LV as a true measure of global cardiovascular efficiency (Asanoi et al., 1989; Tona et al., 2021). The VVC is determined as the ratio of aortic elastance (Ea) and end-systolic elastance (Ees) has recently been recommended for risk stratification for heart failure (Ikonomidis et al., 2019). The parameters, Ea and Ees may be less sensitive in disease, when more sensitive parameters of myocardial longitudinal strain and aortic pulse wave velocity or aortic impedance may be used (Ikonomidis et al., 2019; Alhakak et al., 2021).

In their study, Burchfield et al. (2010) mainly characterized the cardiac function to evaluate the effects of cardiac-restricted low levels TRAF2 overexpression (MHC-TRAF2_LC_) in mice and concluded that TNF is cytoprotective in the heart via TRAF2 mediated activation of NF-kB. We however contend that such models should be studied holistically by evaluating their global cardiovascular function using measurements such as VVC and aortic impedance to determine true global myocardial performance. The goal of this study was to study the same TRAF2 mouse model using cardiac systolic, diastolic, aortic and LV pressure, VVC, and aortic impedance to comprehensively evaluate the baseline global cardiovascular function in comparison to their age matched littermates.

## Methods

### Animals

The generation of MHC-TRAF2_LC_ mice was previously described (Burchfield et al., 2010). Briefly, animals were generated with α-myosin heavy chain promoter targeting murine TRAF2 and characterization of the mice was done using histological analysis at 12 weeks was used to characterize the mice (Burchfield et al., 2010). We used male 8 MHC-TRAF2_LC_ (TRAF2) and 8 littermate controls (CTR) at 2-3 months of age. The diets of both groups of mice consisted of standard commercial chow (2920X Harlan Teklad, Indianapolis, IN, USA) with free access to food and water. All animal protocols were approved by the Institutional Animal Care and Use Committee of Baylor College of Medicine in accordance with the National Institutes of Health Guide for the Care and Use of Laboratory Animals. Anesthesia was induced in mice with 2.5% isoflurane initially, transferred to a heated (37±1°C) electrocardiography board (MouseMonitor S, Indus Instruments, Webster, TX) with isoflurane maintained at 1.5%. The mouse paws were attached to electrodes to measure electrocardiogram (ECG). The experimental setup is and the workflow is shown in Figure 1.

**Figure 1:**
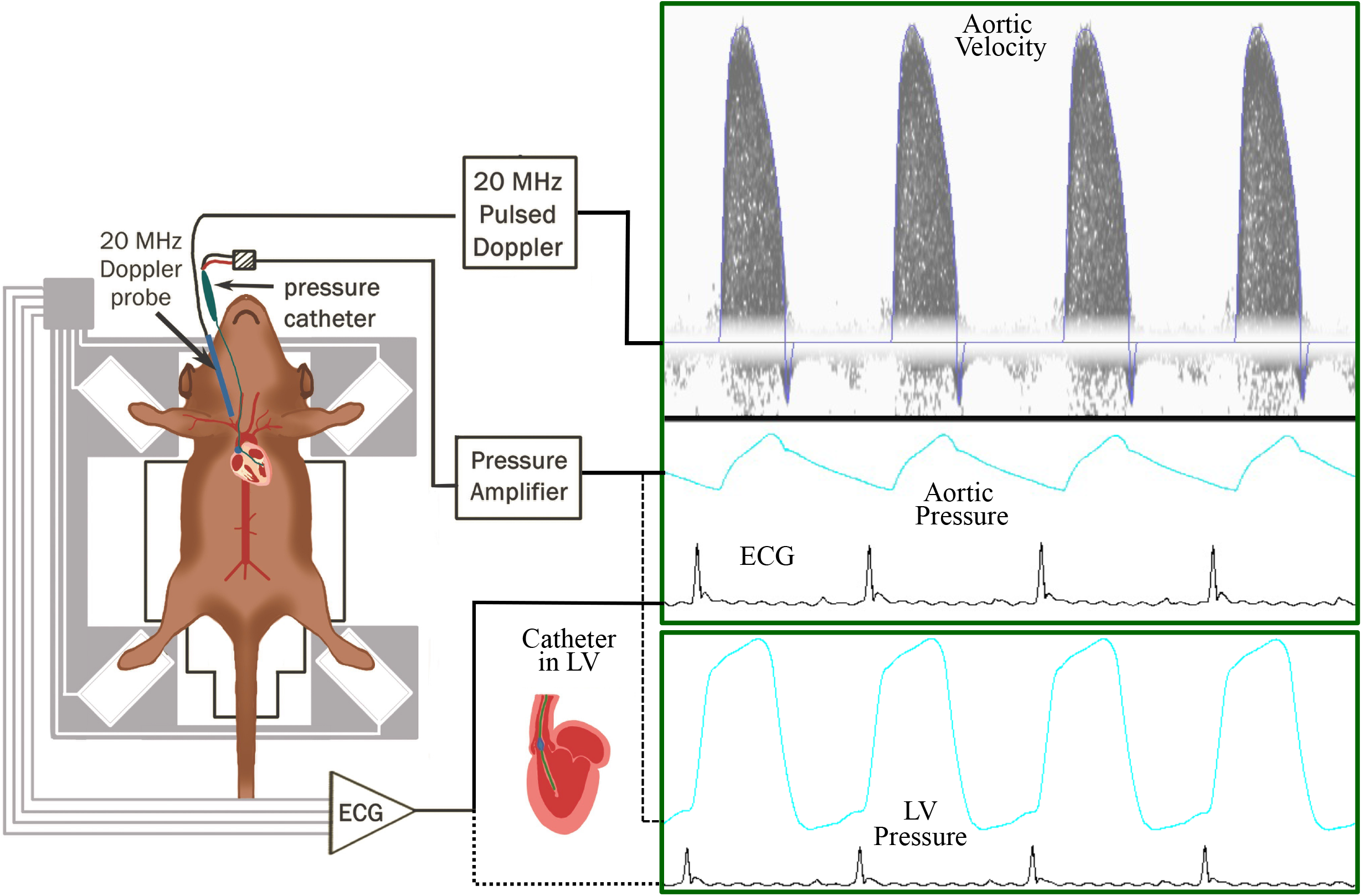
Workflow of the experimental setup to measure aortic blood flow velocity, aortic blood pressure and ECG (electrocardiogram) in TRAF2 and control mice.

### Doppler flow velocity measurements

Cardiac Doppler aortic outflow velocity and mitral inflow signals were measured with 10 MHz Doppler probe with all signals (including blood pressure and ECG) were simultaneously acquired using Doppler Flow Velocity System (DFVS; Indus Instruments, Webster, TX). From the cardiac Doppler signals, we calculated peak (Vp) and mean (Vm) aortic velocities, stroke distance (Sd), ejection time (ET), peak (Ap) and mean (Am) aortic accelerations, Mitral early peak (E) and atrial peak (A) velocities, E/A ratio, deceleration time (DT) isovolumic contraction (IVCT) & relaxation (IVRT) times, and myocardial performance index (MPI - also known as Tei index).

### Blood pressure and flow velocity measurements

Blood pressure measurements were made as previously described (Reddy et al., 2005; Hinton Jr et al., 2016). Briefly, the right carotid artery was isolated and cannulated with a pressure catheter (SPR-1000: Millar Instruments, Inc., Houston, TX) and advanced into the ascending aorta to measure aortic pressure (along with simultaneous measurement aortic blood flow velocity as described below). The aortic pressure and velocity signals were displayed simultaneously, a 2-s segments of both signals were recorded using DFVS system. The catheter was then advanced into the LV. Again, 2-s segments of LV pressure. Systolic (SBP), diastolic (DBP), mean (MBP), pulse pressures (PP), and rate-pressure product (RPP) were calculated from aortic pressure. Peak LV pressure (P_LVP_), indices of contractility (+dP/dt_**max**_) and relaxation (-dP/dt_**max**_), relaxation time constant (tau), and LV end diastolic pressure (LVEDP) were calculated from LV pressure.

### Determination of aortic input impedance

Aortic input impedance was determined using previously reported methods (Reddy et al., 2003, 2005, 2014; Hinton Jr et al., 2016). Briefly, impedance was determined using the simultaneously measured ascending aortic pressure and velocity signals. The pressure and velocity waveforms were processed using discrete Fourier transform to determine aortic input impedance (Zi). Peripheral vascular resistance (Zp – impedance at zero frequency), impedance at first harmonic (Z_1_), characteristic impedance (Zc – average of 2^nd^ to 10^th^ harmonic) were extracted from modulus of aortic impedance (|Zi|). Pulse wave velocity was calculated from |Zi| as Zc/ρ (ρ - density of blood).

### Calculation of parameters to determine VVC

Arterial elastance (Ea) was calculated as ESP/SV (stroke volume, SV=Sd*aortic cross-sectional area), end systolic elastance (Ees) was calculated as ESP/ESV, ventricular-vascular coupling (VVC) was calculated as Ea/Ees, and stroke work (SW) was calculated as ESP*SV (Tona et al., 2021).

### Statistical analyses

All the data are presented as mean ± standard error of the mean (SEM). Statistical analyses were performed via analysis using Student’s T-test on Prism (GraphPad Software; La Jolla, USA).

## Results

The general physiological parameters were measured in TRAF2 mice and compared to CTR mice. Body weight (BW), aortic cross-sectional area (ACA), and heart rate (HR) were unaffected with TRAF2 overexpression (Figure 2A-C). We determined the impact of TRAF2 overexpression on cardiac systolic function and found a significant decrease in peak aortic velocity (Vp), mean aortic velocity (Vm), and stroke distance (Sd) in TRAF2 mice compared to CTR mice (Figure 3A-C). We also found a significant increase in aortic ejection time (ET) (Figure 3D) and significant decreases in both peak (Ap) and mean (Am) acceleration in TRAF2 mice (Figure 3F-G).

**Figure 2:**
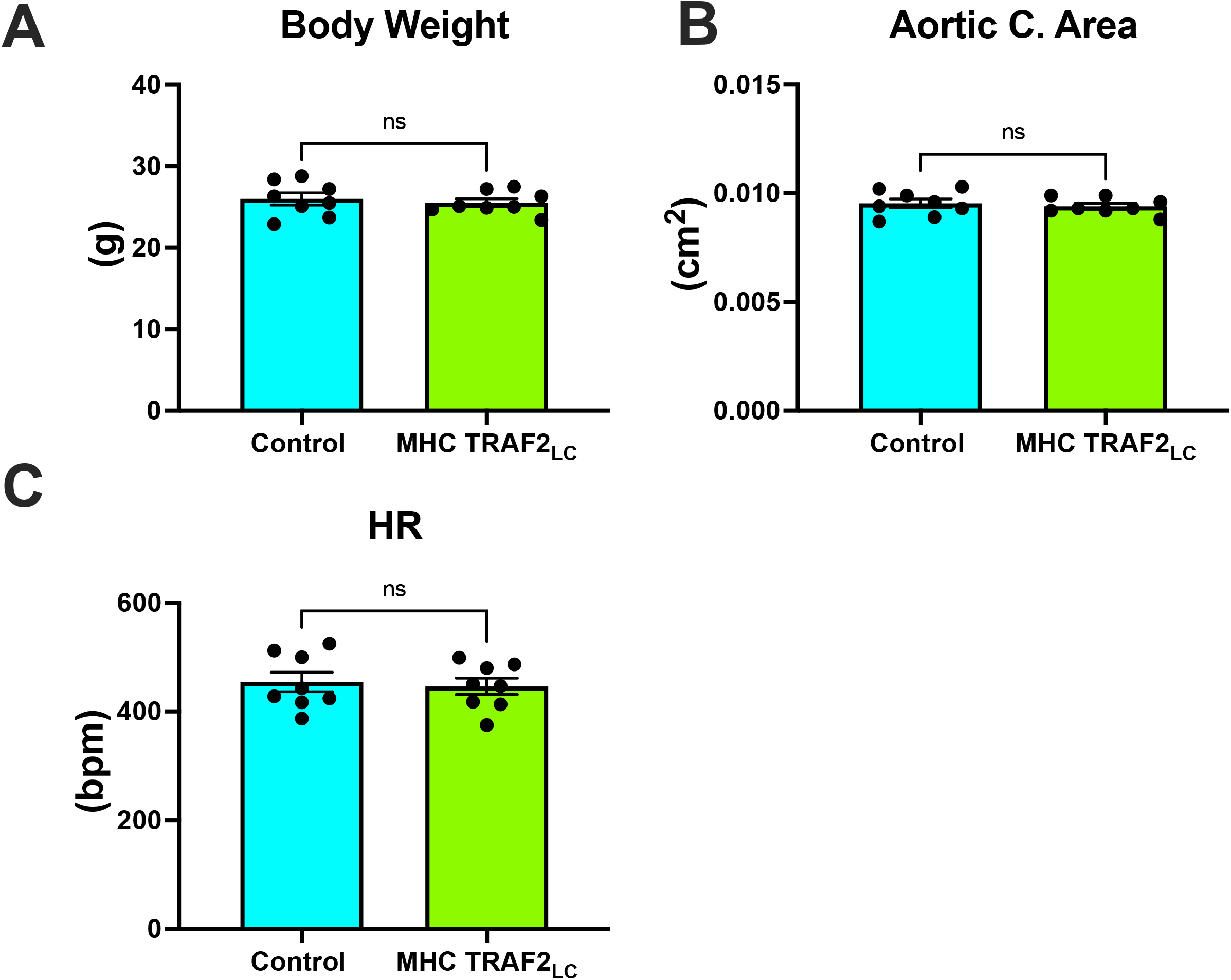
General parameters of male wild type mice and MHC-TRAF2_LC_ mice. (A) Body weight in grams, (B) aortic cross-sectional area, and (C) heart rate (HR) in beats per minute are shown. Data are presented as mean±SEM (n = 8/group). Ns indicates statistically non-significant relationship.

**Figure 3:**
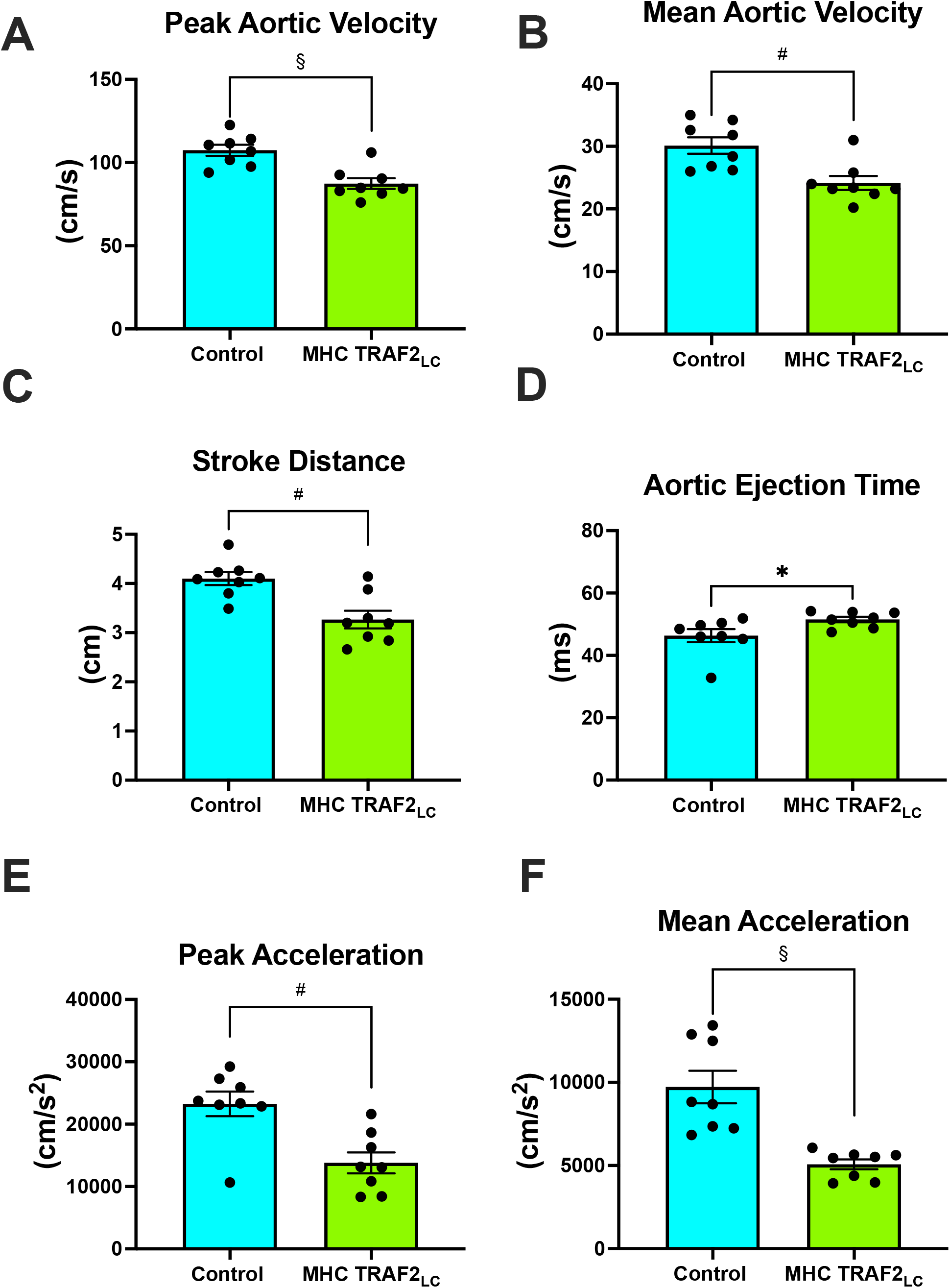
Aortic flow velocity indices of male wild type mice and MHC-TRAF2_LC_ mice. (A) Peak aortic outflow velocity, (B) mean outflow velocity, (C) stroke distance, (D) aortic ejection time, (E) peak aortic outflow acceleration, and (F) mean aortic outflow acceleration. Data are presented as mean±SEM (n = 8/group). *,#, and § represents p <0.05, p < 0.01, and p < 0.001, respectively.

While there were no significant differences in mitral peak early (E) velocity, peak atrial (A) velocity decreased significantly, resulting in the near doubling of Peak E/A ratio (Figure 4A-C). We observed significant decrease in E-deceleration time (DT) and significant increases in both isovolumetric relaxation (IVRT) and contraction time (IVCT) with TRAF2 overexpression (Figure 4D-F). Myocardial performance index (MPI, also known as Tei Index) increased significantly TRAF2 mice compared to CTR mice (Figure 3G).

**Figure 4:**
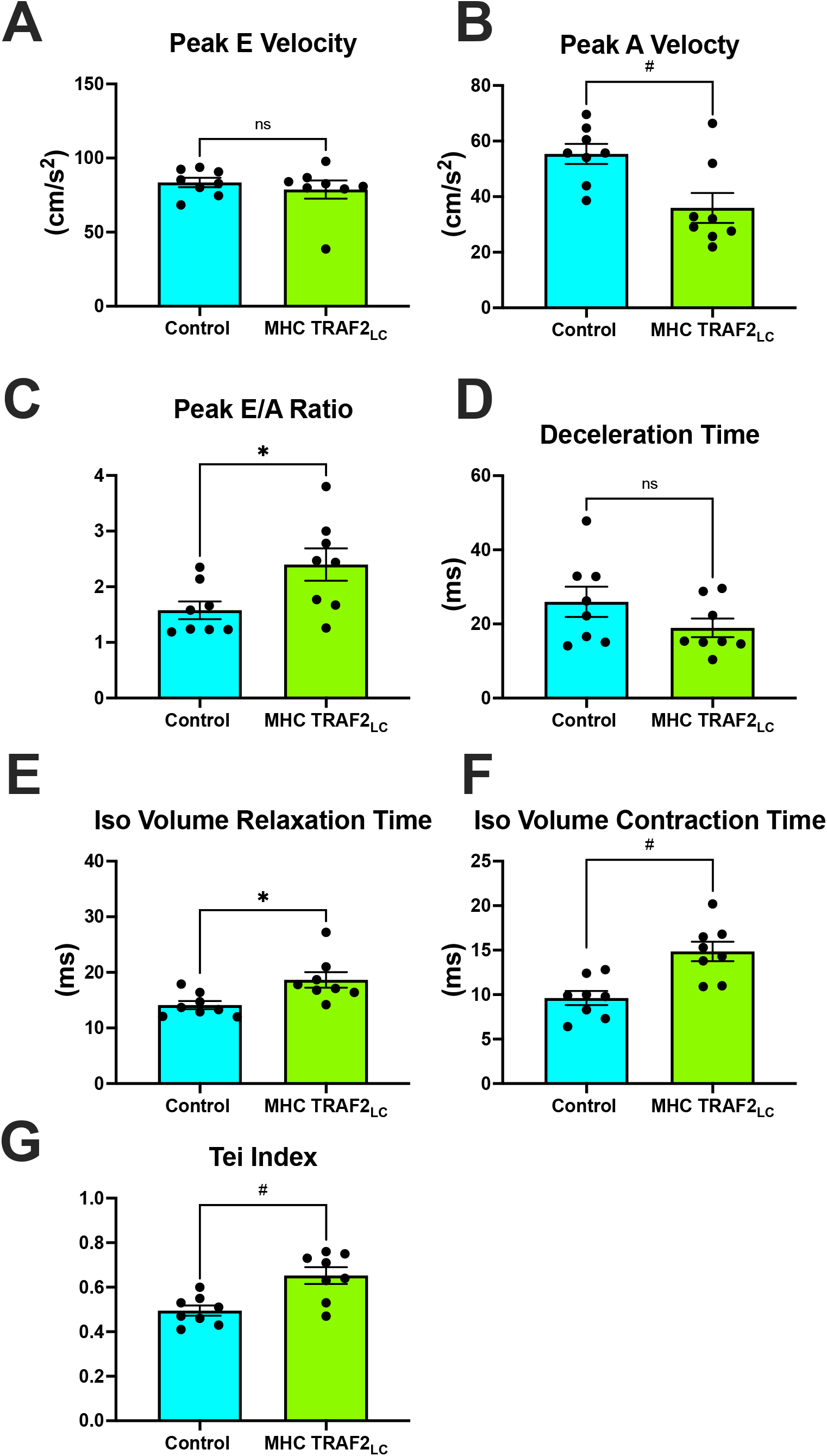
Mitral flow velocity indices of male wild type mice and MHC-TRAF2_LC_ mice. (A) peak mitral-early (E) flow velocity, (B) peak mitral-atrial (A) flow velocity, (C) mitral E/A ratio, (D) E-deceleration time, (E) isovolumic relaxation time, (F) isovolumic contraction time, and (G) myocardial performance index (Tei index), a measure of combined diastolic and systolic function, determined by the sum of isovolumetric relation and contraction time normalized against ejection time. All measurements obtained from cardiac Doppler flow velocity signals in mice. Data are presented as mean±SEM (n = 8/group). * and # represents p <0.05 and p < 0.01, respectively. Ns indicates statistically non-significant relationship.

Systolic blood pressure (SBP) showed a significant decrease with TRAF2 overexpression, while diastolic blood pressure (DBP) showed a slight decrease, but it was not statistically significant (Figure 5A-B). Also, mean blood pressure (MBP), pulse pressure (PP=SBP-DBP), and rate pressure product (RPP=SBP*HR) all decreased significantly decrease in TRAF2 mice compared to the CTR mice (Figure 5C-E).

**Figure 5:**
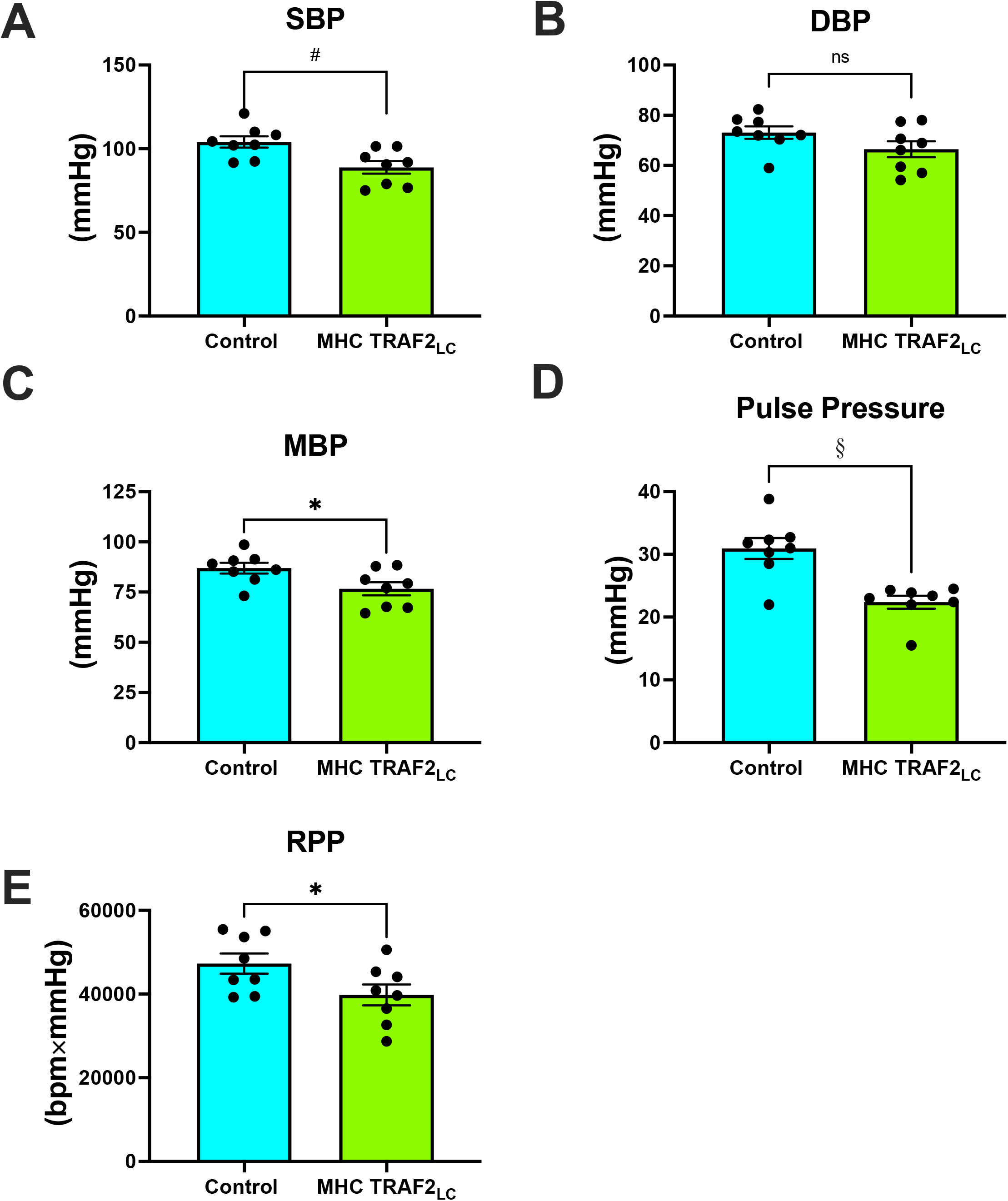
Aortic blood pressure and rate pressure product of male wild type mice and MHC- TRAF2_LC_ mice. (A) Systolic blood pressure (SBP), (B) Diastolic blood pressure (DBP), (C) Mean blood pressure (MBP), (D) Pulse pressure, and (E) Rate pressure product (RPP). Data are presented as mean±SEM (n = 8/group). *,#, and § represents p <0.05, p < 0.01, and p < 0.001, respectively. Ns indicates statistically non-significant relationship.

While peak left ventricular pressure (P_LVP_) was lower in the TRAF2 mice it was not significantly different from CTR mice (Figure 6A), both +dP/dt_max_ and −dP/dt_max_ were significantly lower in TRAF2 mice compared to CTR mice (Figure 6B-C). The relaxation time constant (tau) and LV end diastolic pressure (P_LVED_) trended higher but were not significantly different in TRAF2 overexpression mice compared to CTR mice (Figure 6D-E).

**Figure 6:**
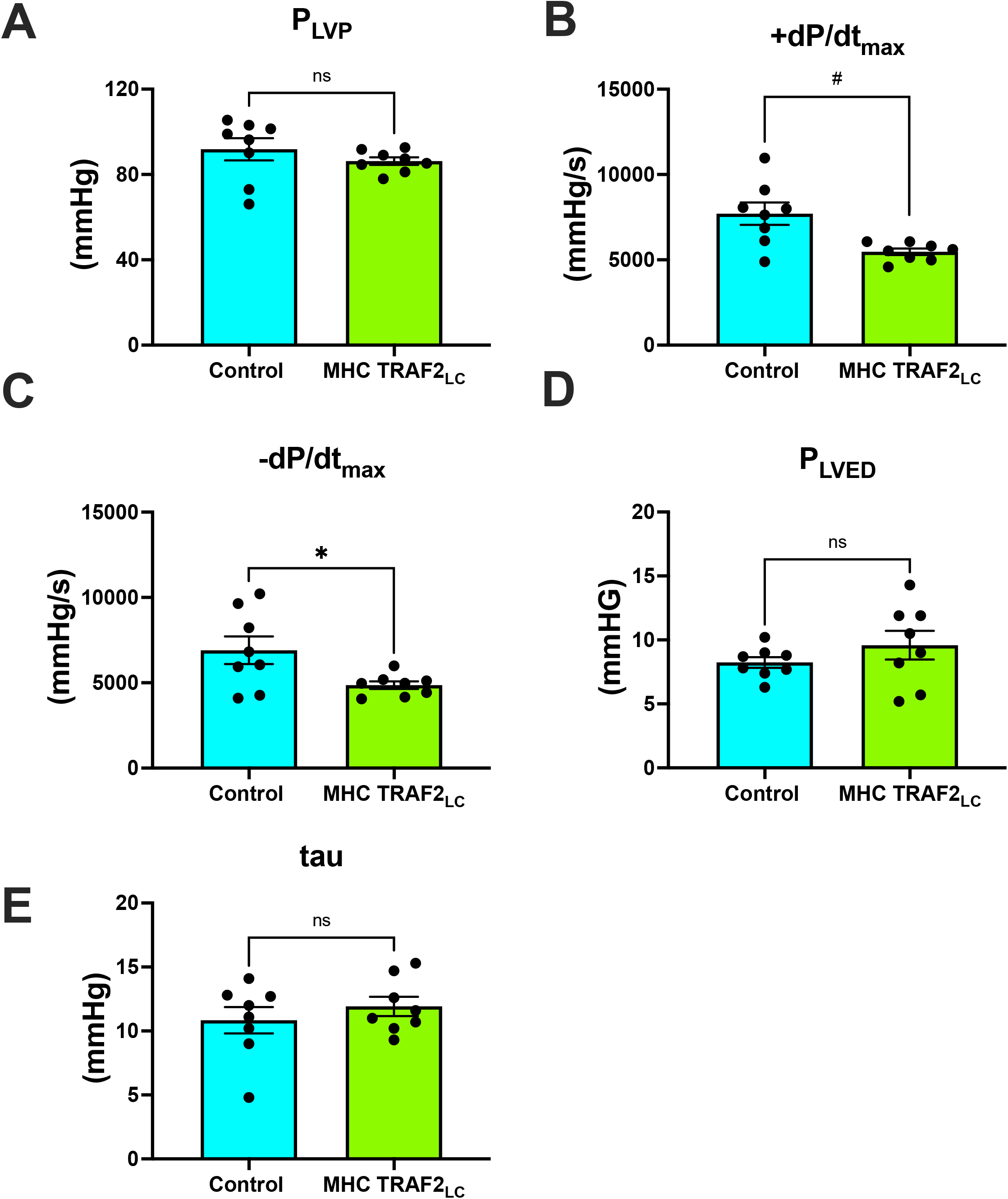
Left ventricular blood pressure parameters of male wild type mice and MHC-TRAF2_LC_ mice. (A) Peak left ventricular pressure (P_LVP_), (B) Maximal contractility (+dP/dt_max_),(C) Maximal relaxation (-dP/dt_max_), (D) left ventricular end diastolic pressure (P_LVED_), and (E) relaxation time constant (tau). Data are presented as mean±SEM (n = 8/group). * and # represents p <0.05 and p < 0.01, respectively. Ns indicates statistically non-significant relationship.

There were no differences in arterial elastance (Ea) but end systolic LV elastance (Ees) was significantly lower in TRAF2 mice, resulting in ventricular-vascular coupling to be significantly higher in TRAF2 mice (Figure 7A-C). Also, stroke work was significantly lower in the TRAF2 mice when compared to CTR mice (Figure 7D)

**Figure 7:**
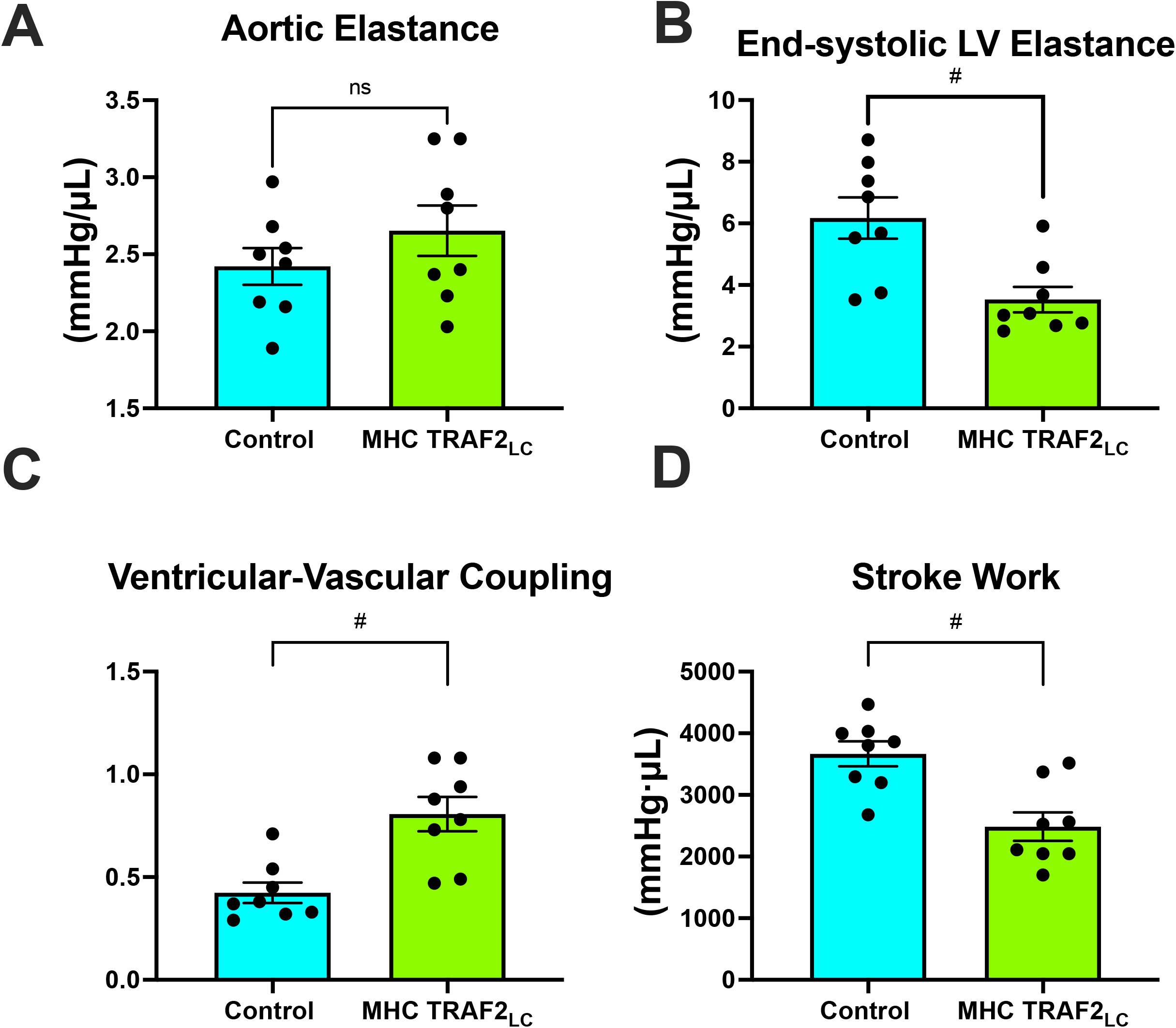
Ventricular-vascular coupling. (A) aortic elastance, (B) end-systolic LV elastance, (C) ventricular-vascular coupling, and (D) stroke work. # represents p < 0.01, and Ns indicates statistically non-significant relationship.

LV afterload was evaluated using aortic impedance. We found no significant differences in any of the impedance parameters, Zp, Z_1_, Zc (or PWVz) between the two groups (Figure 8A-D).

**Figure 8:**
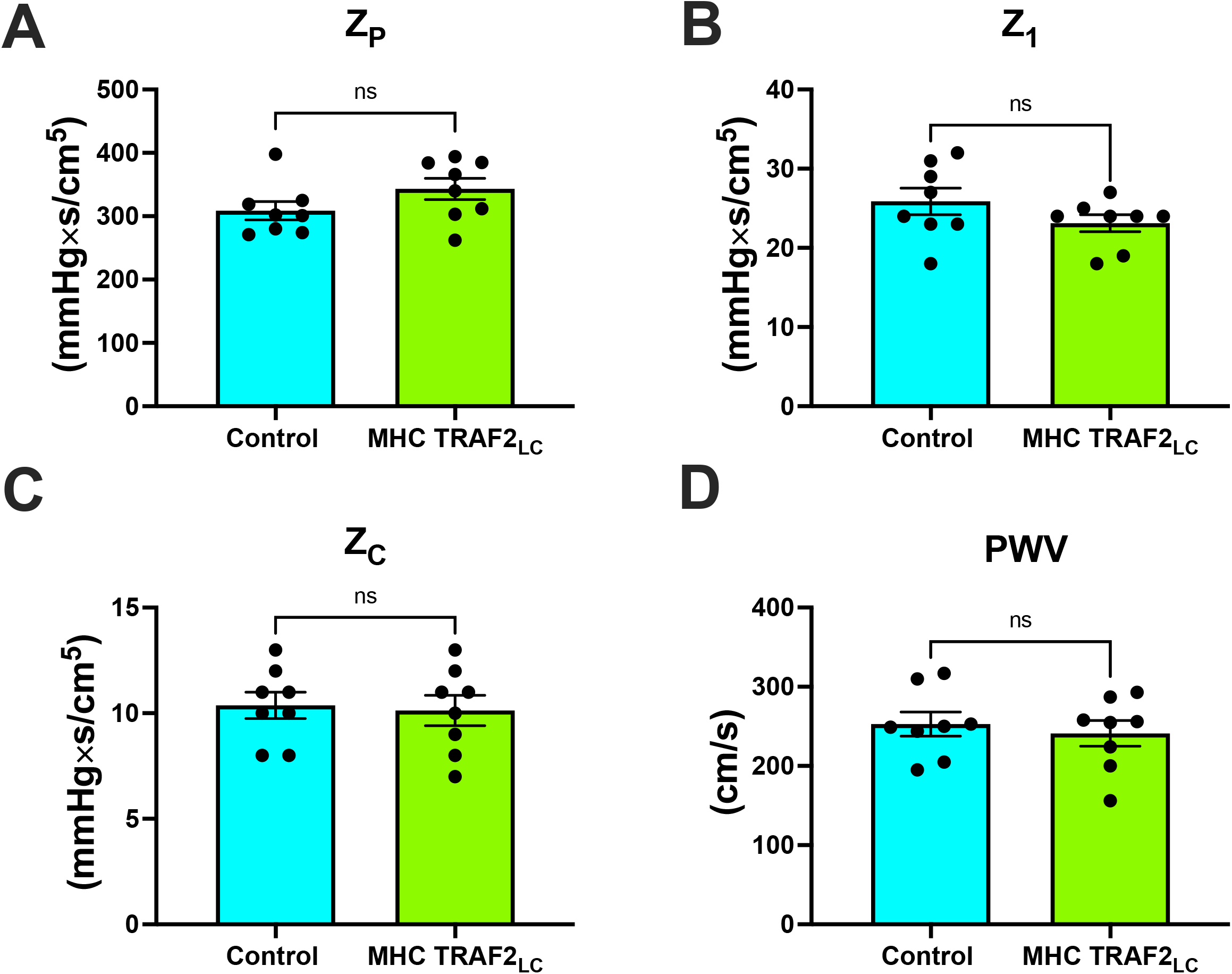
Parameters of aortic impedance in male wild type mice and MHC-TRAF2_LC_ mice. (A) Total peripheral resistance (Z_P_), (B) impedance at first harmonic (Z_1_), (C) characteristic impedance, (Z_C_) (D), and impedance-based pulse wave velocity (PWV). Data are presented as mean±SEM (n = 8/group). Ns indicates statistically non-significant relationship.

## Discussion

Previous study by Burchfied et al. (2010) suggested a TRAF2 mediated cardioprotective function. which was mainly evaluated in the heart. But such models should be studied by evaluating the *in situ* global cardiovascular function. In this study we used invasive measurements of aortic and LV pressure and noninvasive measurements of Doppler mitral inflow and aortic flow velocity and the calculated parameters of VVC and aortic impedance to evaluate the cardiac and aortic hemodynamics in TRAF2 mice and compared to their age matched littermates to determine true global myocardial performance.The general parameters of body weight, aortic cross-sectional area, and heart rate of TRAF2 overexpressing mice were not significantly different from CTR mice (Figure 2).

### Cardiac systolic and diastolic function

We found that the cardiac systolic function, as determined by peak and mean aortic flow velocities, peak and mean aortic accelerations (Figure 3) and +dP/dt_max_ (Figure 4), is diminished in TRAF2 mice compared to the CTR mice which may indicate a mild systolic dysfunction or enhanced systolic reserve. The LV pressure indices of contractility (+dP/dt_max_) and relaxation (-dP/dt_max_) were significantly lower in the TRAF2 mice and followed a similar trend that was observed in the aortic blood pressure and velocity and in agreement with decrease in %FS reported by Burchfield et al. (2010) Again, these diminished contractile and relaxation function in TRAF2 mice are similar to what we observed in the young dwarf *Little* mice (Reddy et al., 2007). We have not challenged these mice to determine systolic reserve capacity, but our group showed that left ventricular developed pressure recovered better in the TRAF2 mice after IR injury and observed that the cytoprotective effects of TRAF2 were mediated by NF-κB activation (Burchfield et al., 2010). But others have reported that suppression of NF-kB with oligonucleotide transcription factor decoys diminishes ischemia related myocardial injury (Tranter et al., 2012) and that cardiac function improved after myocardial infarction with absence of NF-kB p50 unit (Frantz et al., 2006). However, our group reiterated the cytoprotective effects of TRAF2 may have been afforded via crosstalk between canonical and noncanonical NF-kB signaling pathways (Evans et al., 2018). A more recent study noted that TRAF2 maintains myocardial homeostasis through facilitation of cardiac myocyte mitophagy thus preventing inflammation and cell death (Ma et al., 2022). We also found that aortic ejection time were significantly prolonged in the TRAF2 mice (Figure 3D). Obata et al. (2017) reported that longer aortic ejection time is associated with lower blood pressures (Obata et al., 2017), and perhaps the low blood pressures in TRAF2 mice in our study may have caused longer ejection times. Also, longer ET results from longer action potential duration in old animals and may be associated with cellular changes in the mechanics of excitation–contraction coupling (WS Aranow, 2007). Similar cellular changes may be present in TRAF2 that could have caused prolonged ejection time.

The cardiac diastolic function, as defined by E/A ratio was higher in TRAF2 mice than CTR mice (Figure 4C) indicating improved diastolic function. However, E>>A (or E/A>2) and shorter E-deceleration time is associated with moderate decrease in compliance (Oh et al., 2006). The TRAF2 mice had E/A>2 and shorter E-deceleration time (Figure 4D) perhaps indicating a decline in compliance which agrees with the observation by Burchfield et al., (2010) that TRAF2 mice had mildly hypertrophic hearts (Burchfield et al., 2010). Isovolumic contraction time is index of LV contractility and isovolumic relaxation time is an index of LV relaxation, but both are dependent on heart rate (Tei et al., 1997) and other factors. Prolonged IVRT indicates impaired relaxation and prolonged IVCT is associated with reduced EF, both increasing the risk for heart failure (Alhakak et al., 2020). Since IVCT and IVRT are both associated with LV ejection time, the myocardial performance index (MPI or Tei Index) provides an overall cardiac performance (Tei et al., 1997). We found that TRAF2 overexpression resulted a small but significant increase in MPI index (Figure 4G) which may indicate impaired cardiac function like that reported in patients (Harjai et al., 2002).

While TRAF2 overexpression confers cardioprotective effects, the small but significant changes in the systolic and diastolic parameters indicate a modest decrease in baseline function. It remains to be seen if this diminished function deteriorates with age or remains stable over their life span like that in the dwarf *Little* mice which maintain a diminished but stable cardiac function over their life span (Reddy et al., 2007).

### Aortic and LV Pressure

TRAF2 in mice had significantly lower systolic, mean, and pulse pressures compared to CTR mice. The lower blood pressures combined with lower blood velocities in TRAF2 mice make LV afterload appear normal in these mice. Previous studies reported that every 20 or 10 mmHg increase in SBP or DBP, respectively, beyond traditionally healthy values (A Randomized Trial of Intensive versus Standard Blood-Pressure Control, 2017) doubles the chance of major long-term consequences of such as heart failure or stroke (Gebremichael et al., 2019). Since SBP remains a stronger indicator of cardiovascular health than DBP at ages over 50, our findings suggests that the cardioprotective effects of TRAF2 potentially may be conferred across aging (Kjeldsen, 2018). Also, pulse pressure that is ≤25% of systolic pressure is considered too low and typically occurs in aortic stenosis, cardiac tamponade, aortic stenosis, or with blood loss (Homan et al., 2018). Pulse pressure in TRAF2 mice was ≈25% of systolic pressure perhaps indicating modest dysfunction. Given that we measured blood pressure in these mice at young age, studies at older ages may reveal if cardioprotective effects of TRAF2 overexpression are maintained across life span. The rate pressure product (RPP = SBP x heart rate), which is an indirect measurement of the heart oxygen consumption (Nagpal et al., 2007), was significantly lower in TRAF2 mice than CTR mice indicating a lower heart workload.

### Ventricular-Vascular Coupling

A measure of myocardial performance that include interaction with aortic function, the ventricular-vascular coupling (VVC) was significantly higher in the TRAF2 mice indicating a diminished global cardiovascular efficiency. While aortic elastance was unchanged, the higher VVC in TRAF2 mice was mainly due to significantly lower end-systolic LV elastance. Similar observations were made in patients with major adverse cardiovascular events (Tona et al., 2021). Also, it has been shown that decreased LV stroke work index is associated with diminished LV systolic and diastolic function in cardiac intensive care unit patients (Jentzer et al., 2022). Also, we found that stroke work is significantly lower in the in the TRAF2 mice perhaps indicating a diminished cardiac function.

### Aortic Impedance

Defined as the ratio of modulus of pressure to modulus of velocity at several harmonics (0-10) in frequency domain, aortic impedance provides comprehensive description of the afterload experienced by the left ventricle. This includes the pulsatile and the steady components of the hydraulic load compared with when pressure and velocity are considered individually (Reddy et al., 2003; Vlachopoulos et al., 2011; Hinton Jr et al., 2016). We used blood flow velocity instead of volume flow because both blood flow velocity and pressure are independent of body weight or size (Reddy et al., 2003). We found that the peripheral vascular resistance (represented by impedance modulus at 0, Zp), impedance at the first harmonic (Z_1_) which correlates with the strength of the wave reflections from the periphery, characteristic impedance (Zc; average of 2-10 harmonics) which represents the pulsatile component of the aortic pressure-velocity relationship determined by local aortic wall stiffness and diameter and aortic stiffness index, pulse wave velocity (PWV), derived from Zc were not different in TRAF2 mice when compared to CTR mice. The unchanged Zc & PWV_Zc_ seems to conflict with our finding that pulse pressure was lower in TRAF2 mice. While PWV is a direct measure of aortic stiffness, pulse pressure is used as a surrogate stiffness index that also reflects stroke volume (which was also lower in TRAF2 mice) and both measures (stiffness & pulse pressure) are mildly correlated (Waldstein et al., 2008). In another study anti-hypertensive treatments in patients resulted in no changes in aortic PWV but had significant decrease in brachial pulse pressure (Mackenzie et al., 2009).

### Limitations

These observations are made in these mice at young age and the effectiveness of TRAF2 mediated cytoprotection of the heart and cardiovascular system with age needs to be investigated. 2-3 month mice were chosen due to establishing a strong baseline; however, future studies may consider older mice. Female mice may have differential cytoprotection upon TRAF2 activation and therefore may show different effects. Here, we were able to establish both enhanced cardiac reserve but reduced global cardiac function. However, future studies should also consider the mechanism by which this protection is conferred and if this protection is sustained across life span as well as both in male and female mice. Previous studies have found that TRAF2 restores mitophagy, which may be involved with the cytoprotection it provides. While these studies utilized terminal animals, they still provide strong evidence in live model animals. To further understand alterations in echocardiography in live animals, pressure-volume (PV) loops, may be useful in studying cardiac real-time contractile function.

## Conclusions

Previous studies suggest that TRAF2 overexpression may confer cytoprotection of the heart and perhaps afford a potentially important therapeutic target for CVD treatment. However, examination of the global cardiovascular function revealed a diminished cardiac function in the mice with TRAF2 overexpression but there are no differences in the aortic impedance implying no increased afterload. The diminished cardiac function in these mice and the apparent tolerance to ischemic insults may suggest that their cardiovascular reserve may be enhanced. The observations reported by us, and others were from findings in in 2-3 month old mice and perhaps studies need to be conducted at older ages to determine if the cardioprotective effects of TRAF2 overexpression are maintained across life span without significant deterioration of cardiac function.

## Acknowledgement

We like to thank Dr. Douglas L Mann, MD (Professor, Washington University School of Medicine, St. Louis) for providing us the TRAF2 overexpression mice and their littermates.

## Author Contributions

AGM - 3456, KN - 3456, ZV - 3456, HKB - 34, EG - 34, LV - 34, TB - 34, ZE - 34, JA - 34, AM – 34, ES-34, AC-34, JD -34, DS – 34, SD – 34, TTP - 23, JAG - 678, VE - 678, DD - 678, MLE - 7,8, GET - 5678, AOH - 5678, AKR – 12345678

1 Conceived and designed research, 2 Performed experiments and Data collection, 3 Data analysis, 4 Prepared figures, 5 Interpretation of results, 6 Drafted manuscript, 7 Edited and revised manuscript, 8 Approved final version of manuscript.

